# Polyacrylamide bead split-pool method for microbial community analysis

**DOI:** 10.1101/2025.06.26.661670

**Authors:** Niina Smolander, Jasmin Talvitie, Manu Tamminen

## Abstract

Understanding diverse microbial communities is important due to their ecological and medical significance. Bacterial cells are genetically and phenotypically heterogeneous, making their interactions in the communities complex. The heterogeneity and interactions of cells contribute to the formation of specific spatial structures, such as biofilms, and the spread of antibiotic resistance. Here, we describe a novel single-cell approach for studying the cellular heterogeneity and spatial interactions in microbial communities that combines polyacrylamide bead encapsulation of cells and split-pool-barcoding. We demonstrate the method by determining artificially imposed interactions and connecting the taxonomic information in a mock three-species bacterial community with a species-specific genomic target. The method can be utilised for the spatial analysis of microbial communities as well as, once fully optimised for single-cell resolution, linking genetic traits to single cells.

## Introduction

Microbial communities are diverse assemblies of microbes - bacteria, archaea, and microeukaryotes - that co-exist in specific environments and interact with each other and their surroundings. These communities are typically characterized by dense cell populations, high genetic diversity, and complex interactions. Understanding the full complexity of the communities is crucial, given their key roles in host health (e.g., in plants and animals), ecosystem function ^1,2^ and processes such as the spread of antimicrobial resistance ^3,4^. However, much of this complexity remains difficult to resolve, as conventional methods like 16S rRNA gene profiling and metagenomic sequencing lack the resolution needed to capture cell-level interactions and heterogeneity ^5,6^.

Microbes are heterogenous at a cellular level, meaning that even within e.g. a bacterial strain there can be variation between individual cells ^7,8^. Microbial heterogeneity can be categorised into genetic and phenotypic heterogeneity, which in some parts overlap. Genetic heterogeneity is introduced to a community by various mechanisms, including *de novo* mutations, genome rearrangements and horizontal gene transfer (HGT) ^9^. Phenotypic heterogeneity of microbial cells, exhibited for example by variations in gene expression and metabolic activity ^9^, is to some extent driven by genetic heterogeneity and external factors, but is also somewhat stochastic ^10^.

Microbial interactions range from being advantageous for both the interacting cells (mutualism) to being disadvantageous for all (competition). Other potential types of interactions include commensalism, amensalism and predation ^11^. Some microbial interactions are momentary, with no significant impact on the community, while some result in more significant outcomes such as metabolic cross-feeding, HGT and formation of specific spatial structures like biofilms ^11–13^. Beneficial interactions, such metabolic cross-feeding, can lead to physical proximity between cells and ultimately result in formation of specific spatial structures ^14^.

To fully understand complex community structures and the cellular heterogeneity within the communities, methods that can link the bacterial taxonomic information to functional information as well as methods that can assess the physical structure of the community are required. Over the last decade, various methods for studying microbial communities have emerged, such as MaP-Seq^5^ for the study of spatial structure, microfluidic-based BacDrop^15^ and ProBac-seq^16^ for the study of single-cell transcriptomes, as well as microfluidic-based SiC-seq^17^ and Microbe-seq^18^ for single-cell genomics. However, as microfluidic approaches have their limitations, including the requirement of special equipment, other approaches have been developed, such as the use of polyacrylamide beads^19^ and combinatorial barcoding of fixed cells^20^.

Here we describe a method for studying complex microbial communities at a single-cell-resolution. The method combines polyacrylamide beads and modular combinatorial barcoding, also known as split-pool-barcoding, elements previously described by Spencer et al. (2015)^19^, Rosenberg et al. (2017)^21^, Cao et al. (2017)^22^ and Delley & Abate (2021)^23^. In this method bacterial cells are first trapped to irreversible polyacrylamide beads, and DNA of interest is amplified using emulsion PCR. The amplicons are then barcoded using split-pool barcoding, detached from the beads, amplified and finally sequenced using next generation sequencing. The described method requires no specialized laboratory infrastructure and is easy to set up in any laboratory setting.

## Methods

### Polyacrylamide beads

An acrylamide mixture was made by combining 10 µL of bacterial cells (suspended in 1xPBS), 52 µL of 1xTE buffer, 5 µL of each 100 µM acrydited forward primer (/5ACryd/ATGC/ideoxyU/GTGCCAGCMGCCGCGGTAA, /5ACryd/ATGC/ideoxyU/TTATAACTTCCAAAGGGCTGAC, /5ACryd/ATGC/ideoxyU/GCCGTCAACTACTATCTTGTAA; Integrated DNA Technologies (IDT)), 3 µL of 10% ammonium persulfate and 20 µL of 30% acrylamide/bisacrylamide suspension (37,5/1 ratio, Sigma-Aldrich). A RAN-TEMED solution was made by adding 0.5 µL of >99 % Tetramethyl ethylenediamine (Sigma-Aldrich) to 100 µL of HFE7500 + 20g of 5 weight-% 008-FluoroSurfactant RAN oil (RAN Biotechnologies). The acrylamide mixture was added to the RAN-TEMED solution and emulsified using a 150 µL pipette by pipetting in and out 20-30 times. 100 µL of mineral oil (Sigma-Aldrich) was added on top without mixing, and the emulsion was incubated at 65°C for 16-18 h. Two set of polyacrylamide beads were made: one containing *Comamonas testosteroni* and *Brevundimonas bullata* cells, the other one containing *Agrobacterium tumefaciens* cells. The cells were provided by Microbial Domain Biological Resource Centre HAMBI (Biodiversity Collections Research Infrastructure (HUBCRI), Helsinki Institute of Life Science (HiLIFE)).

After the incubation, the beads were extracted and cleaned up. First, the bead-oil mixture was spun for 5 s, and the top and bottom oil phases were removed. Next, 50 µL of 97% perfluoro-1-octanol (Sigma-Aldrich) and 100 µL of 1xTE buffer were added to the mixture and mixed by tapping. The sample was spun for 5 s and the bottom phase was discarded. To harvest the beads, 500 µL of 1xTE buffer was added, and the mixture spun for 5-10 s. Most of the top (bead) phase was collected to a new tube. To wash the beads, 1 mL of 1xTE buffer was added and the samples were centrifuged for 1 min at 13,000 rcf. The supernatant was removed, and the washing step was repeated. After the second wash, the beads were resuspended in 500 µL of 1xTE buffer. The beads were vortexed and 70µm size filtered with a cell strainer.

### Multiplex emulsion PCR

Three targets were amplified in the multiplex emulsion PCR: 16S rRNA gene region V4, 299 bp long target specific to *A. tumefaciens* and 283 bp long target specific to *C. testosteroni*. First, the two sets of polyacrylamide beads with different bacterial cells were combined. For one reaction, 20 µL of 5x HF buffer (NEB), 2 µL of 50mM MgCl_2_ (NEB), 2.5 µL of 10mM dNTPs (NEB), 8 µL of 2,000 units/ml Phusion® Hot Start Flex DNA Polymerase (NEB), 2.5 µL of each 40 µM forward primer (CAGCMGCCGCGGTAATWC, TTATAACTTCCAAAGGGCTGAC, GCCGTCAACTACTATCTTGTAA; IDT), 2.5 µL of each 40 µM reverse overhang primer (ACCACGCTCCAATTAAGCGGGGACTACHVGGGTWTCTAAT, ACCACGCTCCAATTAAGCGGGACGAGATAATAATTGAGCGC, ACCACGCTCCAATTAAGCGGTGCCTATATCGTTCCAGACGG; IDT) and 5 µL of sterile H_2_O was added. 52.5 µL of the reaction mix was combined with 47.5 µL of polyacrylamide beads, three 4 mm glass beads (Supelco®) and 900 µL of a solution containing 4% ABIL EM 90 (Evonik) and 0.05% Triton X-100 (Sigma-Aldrich) in mineral oil. The samples were emulsified by vortexing for 1 min. Each reaction was divided into sixteen 60 µL aliquots. The amplification programme was 30 s initial denaturation at 94°C followed by 20 cycles of denaturing at 94°C for 10 s, annealing at 55°C for 30 s, and extension at 72°C for 12 s. The final extension was carried out at 72°C for 5 min.

After the PCR, the aliquots were pooled to a 2mL tube, and the beads were extracted. First, 1 mL of water-saturated diethyl ether was added, and the samples were vortexed and centrifuged for 1 min at 13,000 rcf. The top phase was discarded, and the step was repeated. Next, 1 mL of water-saturated diethyl acetate was added to each sample. The samples were mixed by tapping and centrifuged as before. The top phase was discarded. Next, 1 mL of diethyl ether was added. The samples were mixed by tapping gently and the top phase was discarded. The diethyl ether wash step was repeated, and the ether solvent was allowed to evaporate from the samples for at least 3 mins. The samples were then transferred into new 1.5 mL tubes and the previous 2 mL tubes were rinsed with 500 µL 1xTE buffer to collect all the material. The samples were centrifuged for 1 min at 13,000 rcf, the supernatant was discarded, and the samples were resuspended to 200 µl with 1xTE buffer. The samples were combined with 200µl of 0.2M NaOH, vortexed and incubated for 10 min. The mixture was then vortexed and centrifuged for 1 min at 13,000 rcf. The supernatant was removed, the beads were washed twice with 200 µL of 1xTE buffer and resuspended to 500 µl of 1xTE buffer.

### Split-pool barcoding

Prior to the split-pool barcoding, three linker-barcode plates and blocking oligos were prepared (see Supplementary Data 1 for sequences). First, the blocking oligos, three barcode 96-well plates and three linker oligos were diluted to a concentration of 10 µM. 5 µL of linker 1 was transferred to each well of a 96-well plate. Subsequently, 5 µL of barcode from each well of the 10 µM barcode 1 plate was added to respective wells in the linker 1 plate. To hybridise the barcodes to the linkers, the linker-barcode plate was heated to 95°C and then cooled down to 20°C at a rate of -0.1°C/s. The same steps were repeated for linker-barcode plates 2 and 3.

For the split-pool barcoding of the beads, a ligation mix was prepared on ice by combining 265 µL of 10x T4 Ligase buffer (NEB), 530 µL of polyacrylamide beads, 742 µL of sterile H_2_O and 53 µL of T4 Ligase (NEB). 15 µL of the ligation mix was transferred to each well of the linker-barcode plate 1. The linker-barcode-ligation mix plate was incubated for 30 mins at 37°C with 300 rpm shaking and then heated to 65°C for 20 mins. Unhybridized linkers were blocked with the blocking oligo. Blocking mixture contained per reaction, 1.25 µL of 10x T4 Ligase buffer, 0.6 µL of the blocking oligo 1 and 3.15 µL of sterile H_2_O. 5 µl of blocking mixture was added to each well of the linker-barcode-ligation mix plate and the plate was incubated for 30 mins at 37°C with 300 rpm shaking. After the incubation, the wells were pooled, and the material was washed first twice with 0.1% SDS and then three times with 1xTE buffer. After washes the beads were resuspended to 530µl with 1xTE buffer.

The protocol was repeated twice. In the second and third round of the split-pool barcoding, linker-barcode plates 2 and 3 were used, respectively, and the blocking mix contained per reaction 0.8 µL of respective blocking oligo, 1.35 µL 10x T4 Ligase buffer and 2.85 µL sterile H_2_O. Finally, the beads were treated with NaOH as previously and washed with and resuspended in 1xTE buffer.

### DNA releasing and amplification

The barcoded DNA was detached from the beads by combining the beads with 1µl of 1,000 units/mL USER enzyme per 10µl of sample and incubating at 37°C for 1 h. After incubation, the sample was vortexed for 1 min, frozen for at least 1 h, vortexed for 2 min and incubated at 37°C for 2 h in 300 rpm shaking. Finally, the sample was centrifuged for 1 min at 13,000 rcf and the supernatant was collected.

The barcoded 16S rRNA gene region V4 and the barcoded target specific to *A. tumefaciens* were separately amplified in two subsequent PCRs. In the first PCR, the DNA was just amplified using the same forward primers as in the emulsion PCR (CAGCMGCCGCGGTAATWC and TTATAACTTCCAAAGGGCTGAC, 16S rRNA gene and *A. tumefaciens*-specific, respectively; IDT) and a barcode-end-targeting reverse primer (TCTCCAAATGGGTCATGATC; IDT). In the second PCR, purified DNA (Monarch® PCR & DNA Cleanup Kit (NEB)) from the first PCR was amplified and frameshifts and adapter sequences were introduced using an overhang forward (GTCTCGTGGGCTCGGAGATGTGTATAAGAGACAGNCAGCAGCCGCGGTAATAC and GTCTCGTGGGCTCGGAGATGTGTATAAGAGACAGNTTATAACTTCCAAAGGGCT GAC, 16S rRNA gene and *A. tumefaciens*-specific, respectively; IDT) and reverse (TCGTCGGCAGCGTCAGATGTGTATAAGAGACAGNCATCTTCTCCAAATGGGTCA TGAT; IDT) primer mixes that target the same sites as the primers in the first PCR. The primer mixes consisted of four primers with 0-3 N nucleotides in the middle. Per reaction, the PCR reactions consisted of 4µl of 5x Phusion HF buffer (NEB), 0.4 µL of 10mM dNTP Solution Mix (NEB), 1 µL of each 10µM primer / primer mix, 0.2 µL of 2,000 units/mL Phusion® Hot Start Flex DNA Polymerase (NEB) and 13.4 µl of template DNA (1st PCR) or 10µl of template DNA and 3.4µl of H_2_O (2nd PCR). The amplification programme in the first PCR was 30 s initial denaturation at 98°C followed by 20 cycles of denaturing at 98°C for 10 s, annealing at 61°C for 30 s, and extension at 72°C for 15 s. The final extension was carried out at 72°C for 5 min. The second PCR followed the same protocol apart from the annealing was carried out at 63°C +0.5°C/cycle for 10 cycles. After the second PCR, the amplicons were run on Invitrogen E-Gel EX (2% agarose) for 10 min and the DNA sized approximately 485-515 bp was excised and purified with Monarch® PCR & DNA Cleanup Kit (NEB).

### Library preparation and sequencing

The barcoded 16S rRNA gene and *A. tumefaciens*-specific PCR amplicons were indexed for sequencing separately. Per reaction, the PCR reactions consisted of 4 µL of 5x Phusion HF buffer (NEB), 0.4 µL of 10mM dNTP Solution Mix (NEB), 1 µL of 10 µM forward (i5) and reverse (i7) primers each, 0.2 µL of 2,000 units/mL Phusion® Hot Start Flex DNA Polymerase (NEB), 11.4 µL of sterile H_2_O and 2µl of the template DNA. The indexing programme was 30 s initial denaturation at 98°C followed by 8 cycles of denaturing at 98°C for 10 s, annealing at 55°C for 20 s, and extension at 72°C for 20 s. The final extension was carried out at 72°C for 5 min. The amplicons were run on Invitrogen E-Gel EX (2% agarose) for 10 min and the DNA sized approximately 558 bp was excised and purified with Monarch® PCR & DNA Cleanup Kit (NEB). The two purified samples were combined in equimolar concentration to a 4 nM library. The DNA was sequenced with Illumina MiSeqTM system using the MiSeq® Reagent Nano Kit v2 (500 cycles) at the Finnish Functional Genomics Centre (Turku, Finland).

### Bioinformatics and data analysis

The adapter sequences were trimmed by the Illumina FASTQ file generation pipeline, and the quality of the reads was evaluated using FastQC v0.12.1^24^. The reads from the sequencing were converted from FASTQ to FASTA format using SeqKit^25^ and processed using R v.4.4.3^26^ and RStudio^27^. Only the reads with a correctly formed barcode sequence were kept for the analysis (see Figure 1 for reference): (1) the expected sequence was found in the non-variable regions between the barcode 3 (BC3) and BC2 variable regions, BC2 and BC1 variable regions and the BC1 variable region and the target amplicon; (2) the three variable regions were one of the 96 possible barcode sequences; (3) the unique molecular identifier (UMI) situated in the first barcode oligo was 10 nucleotides long.

**Figure 1.**
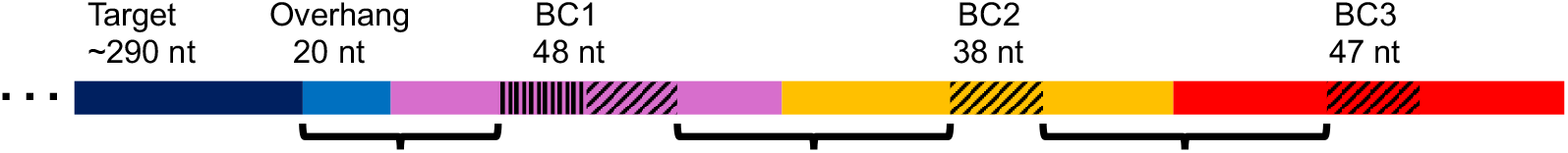
Split-pool barcode structure. From left to right: target DNA attached to the bead (dark blue), overhang region (blue), barcode 1 (BC1, purple), BC2 (orange) and BC3 (red). The 10-nt UMI region indicated with vertical lines and the 8-nt variable regions in each BC with diagonal lines. Curly brackets indicate the non-variable regions used for the filtering of the barcodes.

The sequences outside of the barcode sequence were mapped against the 16S rRNA gene sequences of *Agrobacterium tumefaciens, Brevundimonas bullata* and *Comamonas testosteroni* and the expected amplicon sequence of the *A. tumefaciens*-specific target using the R package blaster v.1.0.7^28^ with minimum identity of 95%. The mapping was performed separately for the reads from the barcoded 16S rRNA gene and *A. tumefaciens*-specific target samples. Reads that could not be assigned or with multiple hits were filtered out.

The three 8-nucleotide barcode sequences were concatenated into one full barcode. The (UMI) sequence included in the first barcode was used for filtering. The number of times each UMI was associated with each full barcode and each target sequence was counted and only the barcode and the target sequence with clearly the highest count (≥ 70% of the counts) for each UMI was kept. Hence, the UMIs with no clear association to just one barcode or target sequence were filtered out. The results were visualised using ggplot2 ^29^.

## Results

Figure 2 summarises the key steps of the polyacrylamide bead split-pool barcoding protocol described in this manuscript. An excess of bacterial cells was used in encapsulation to ensure that all polyacrylamide beads had at least a single bacterial cell. Cells of *B. bullata* and *C. testosteroni* were co-encapsulated to mimic a physical interaction between these cell types, while *A. tumefaciens* was encapsulated separately to mimic the lack of interaction between other cells.

**Figure 2.**
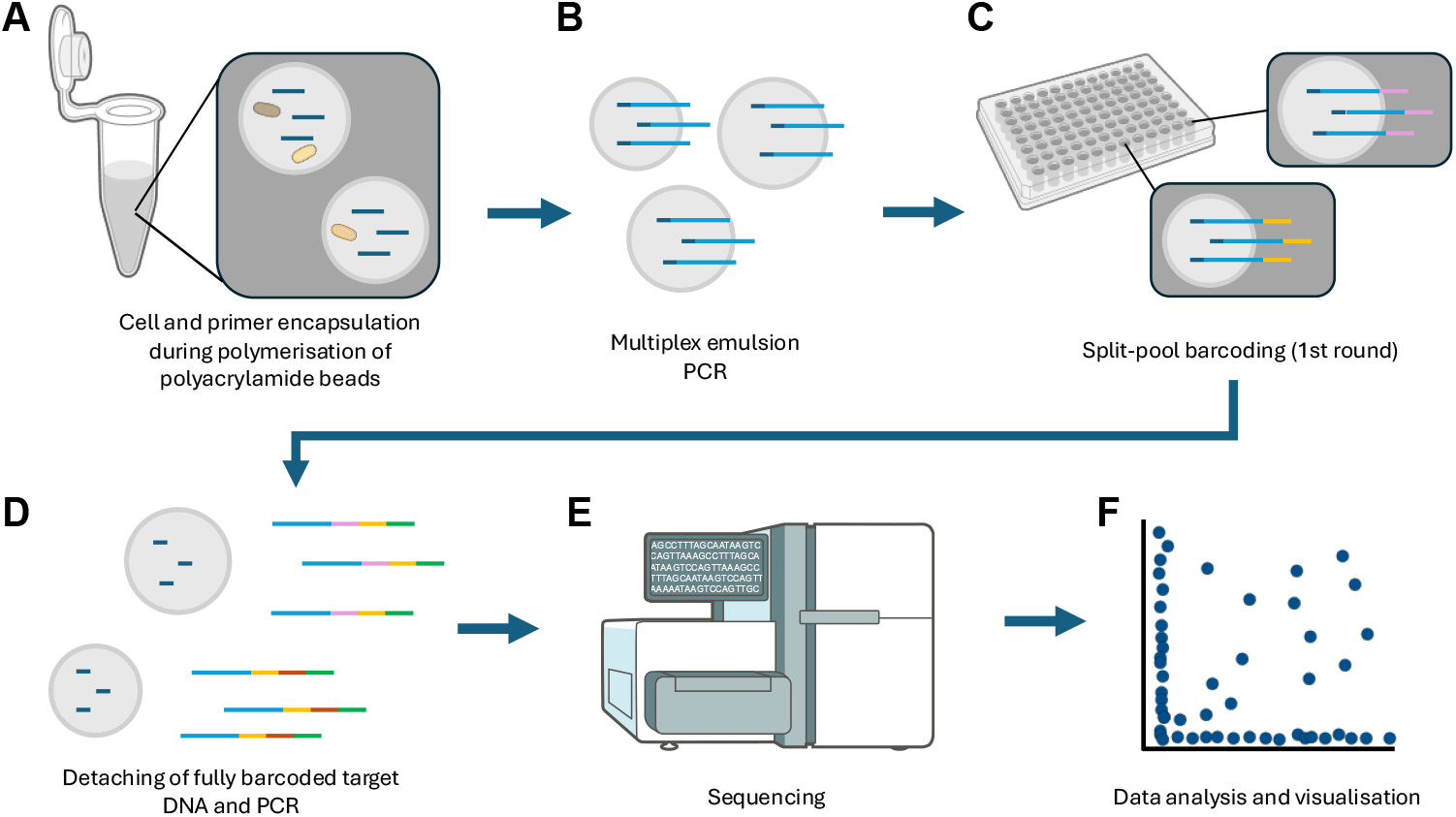
Polyacrylamide split-pool barcoding method overview. (a) The bacterial cells were encapsulated into polyacrylamide beads by suspending the cells into acrylamide-TEMED-oil suspension, mixing the suspension into an oil emulsion by pipetting, and allowing the emulsion droplets to polymerize. Acrydited primers were immobilised during the polymerisation of the beads. After polymerization, the beads were extracted from the oil emulsion and washed. (b) The DNA of interest was amplified in a multiplex emulsion PCR, the amplicons attaching to the beads via the acrydited primers. (c) The amplicons were split-pool barcoded, (d) detached from the polyacrylamide beads and the barcoded DNA was amplified. (e) The DNA was sequenced with the Illumina MiSeq^TM^ system and (f) the sequencing reads were analysed. Figure created using illustrations from NIAID NIH BIOART.

### Polyacrylamide bead matrix remained intact throughout the split-pool protocol

Throughout the protocol – after the polymerization and extraction of the polyacrylamide beads, after the emulsion PCR and during the split-pool barcoding – the beads were examined microscopically. The bead size varied between 20-100 µm, the mean size being approx. 60-70 µm. It was also confirmed that the bacterial cells remained encapsulated throughout the protocol and that the beads remained intact and did not aggregate. Some loss of beads was encountered during the protocol. (Figure 3).

**Figure 3.**
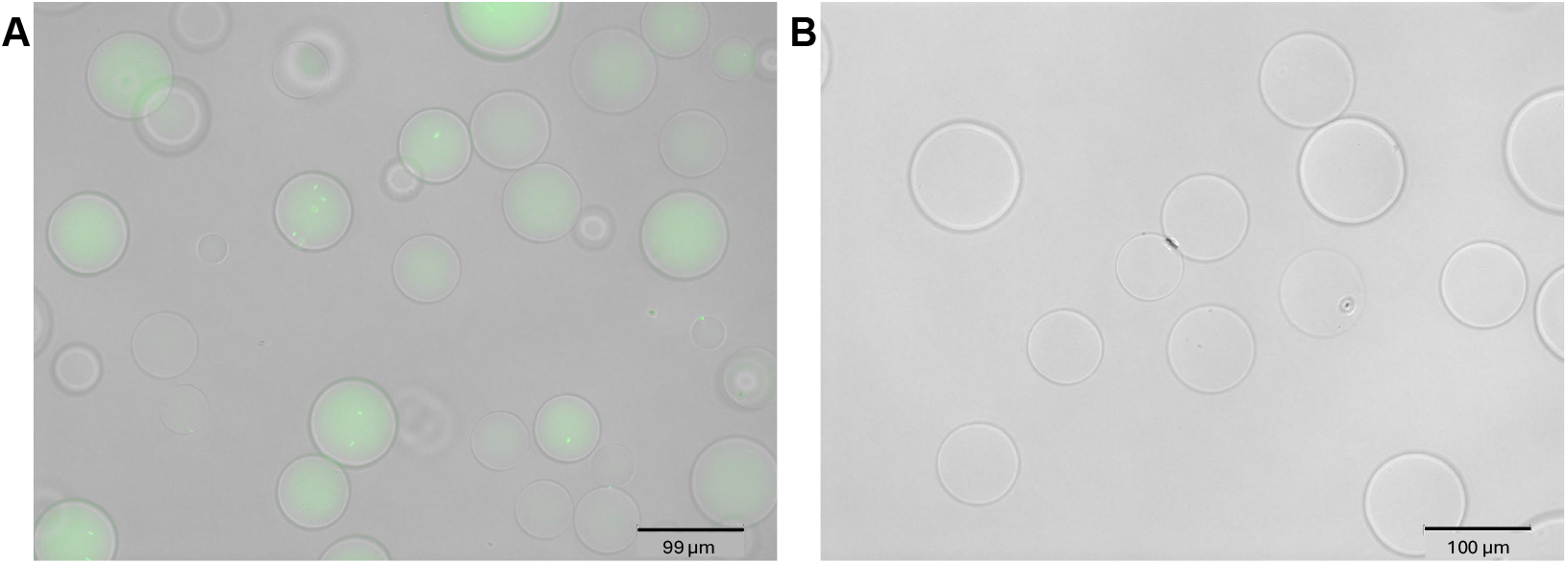
Microscope images of the polyacrylamide beads (a) after the emulsion PCR, DNA stained with SYBR® Safe DNA Gel Stain (Invitrogen) (b) after the split-pool barcoding.

### The barcode sequence quality

In the multiplex emulsion PCR, prior to the split-pool barcoding, three barcoding targets were amplified: 16S rRNA gene region V4, a 299 bp long genomic target specific to *A. tumefaciens* and a 283 bp long genomic target specific to *C. testosteroni*. Due to issues with the *C. testosteroni*-specific target, it was not included in the post-barcoding PCRs. After the split-pool barcoding, the barcoded DNA from approximately 1000 beads was detached, and the DNA was amplified in two subsequent PCRs and indexed for sequencing (Figure 4).

**Figure 4.**
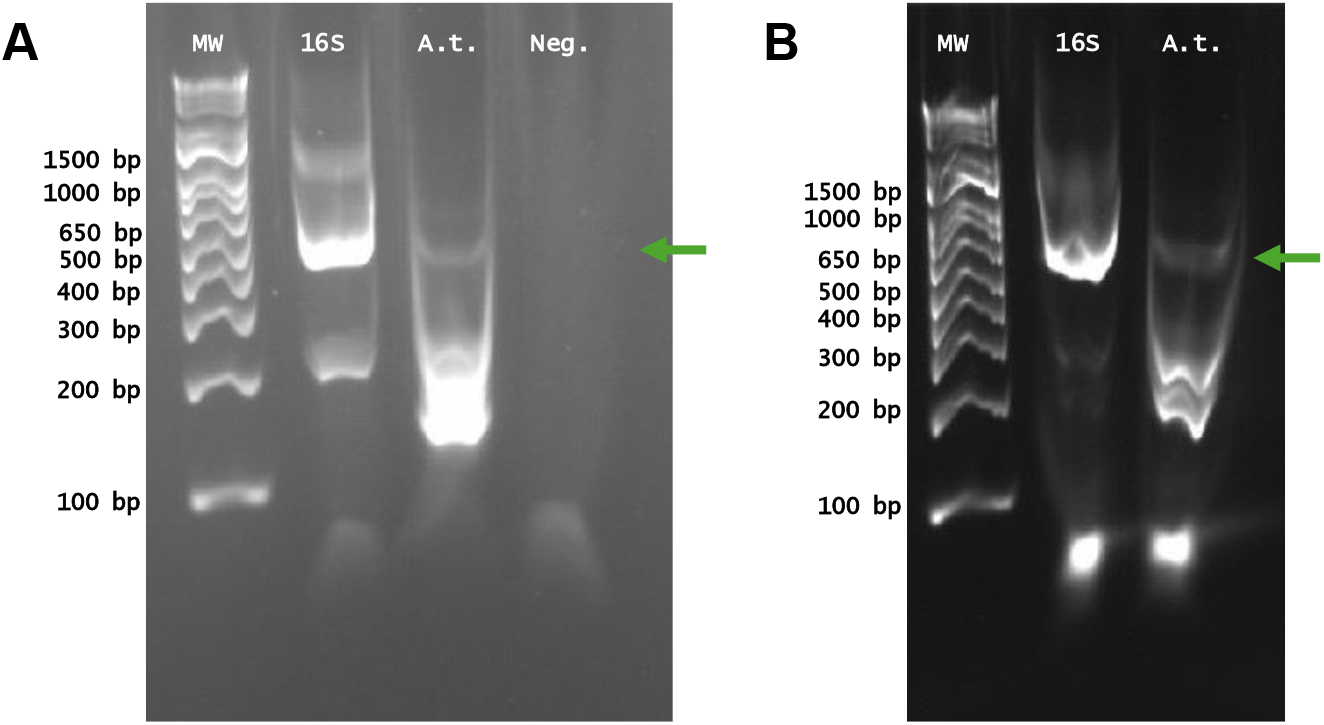
Gel electrophoresis images of barcoded 16S rRNA gene and A. tumefaciens-specific target (A.t.) after (a) the second PCR (frameshift PCR) and (b) indexing PCR. The correctly sized 16S rRNA gene and A. tumefaciens genomic target (A.t.) bands excised from the gel are indicated with green arrow. Negative control (Neg.) included only in the frameshift PCR. MW = molecular weight (1Kb Plus DNA Ladder). Gel run 48V, 10 min on 2% agarose E-Gel^TM^ EX Agarose Gel.

Altogether, the sequencing returned 739,421 barcode-end reads (Supplementary Data 2) of which 255,231 contained a barcode sequence that followed the expected structure. Majority of the barcoded 16S rRNA gene reads were correctly formed (174,307 out of 242,826 reads), while the opposite was the case for the *A. tumefaciens*-specific reads (80,924 out of 496,595 reads). There were 96, 95 and 96 unique barcode 1, barcode 2 and barcode 3 sequences, respectively, meaning nearly all of the potential 96 barcodes from each split-pool barcoding round were sampled.

Of those reads with a complete and correctly formed barcode region, 165,739 could reliably be assigned a bacterial strain and 66,480 the *A. tumefaciens*-specific target. When the three variable barcode regions were concatenated, altogether there were 12,399 unique barcode sequences. The sampling range of the barcodes was 1–26,446, the median and mean sampling counts being 1 and 19 respectively.

A 10 nucleotide long unique molecular identifier (UMI) sequence was included in the first barcode oligo. In those reads where a bacterial strain was assigned, altogether 7,982 unique UMI sequences where found. The sampling range of the UMIs was 1–192, the median and mean sampling counts being 26 and 29 respectively. After the filtering based on the UMIs was performed, there were 7,940 unique UMIs and 1,583 unique barcodes.

### Barcode distribution between bacterial strains

Of the complete and correctly formed barcodes, 262 were exclusively associated with *A. tumefaciens*, 258 with *B. bullata*, 538 *C. testosteroni* and 118 with the *A. tumefaciens*-specific genomic target (Figure 5). Some of the barcodes were shared between the targets with the greatest number of associations being between *B. bullata* and *C. testosteroni*, as was expected based on their deliberate co-encapsulation to mimic a physical interaction between these cells (Figure 5). Additionally, the *A. tumefaciens*-specific genomic target shared more barcodes with the *A. tumefaciens* 16S rRNA gene than with *B. bullata* or *C. testosteroni*, as expected. 74 barcodes were found in association with any three of the targets and 6 with all four targets. Ideally, in the current set up, the number of these barcodes should have been minimal.

**Figure 5.**
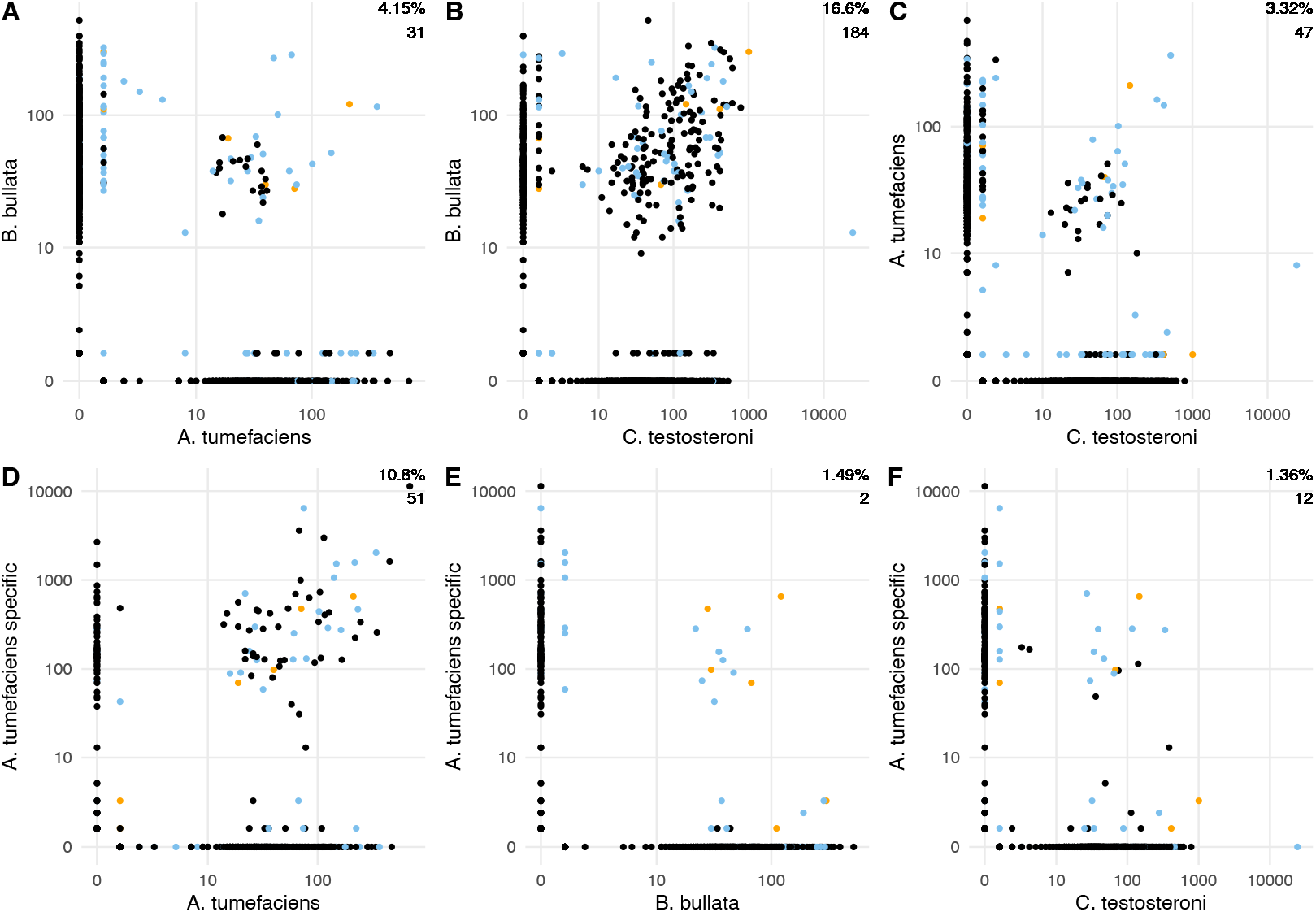
Number of times each barcode was associated with the targets. Each point represents a barcode and its position the number of times it was associated with (a) A. tumefaciens or B. bullata, (b) C. testosteroni or B. bullata, (c) C. testosteroni or A. tumefaciens, (d) A. tumefaciens or A. tumefaciens-specific target, (e) B. bullata or A. tumefaciens-specific target and (f) C. testosteroni or A. tumefaciens-specific target. Blue points represent barcodes that were associated with three targets and orange points barcodes that were associated with all four targets. Percentages represent the proportion of barcodes that were ≥ 10 for both respective targets. The values under the percentages are the number of barcodes associated exclusively just with both respective targets (i.e. barcodes associated with three or four targets are not included in the values). X and Y axes are on pseudo-logarithmic scale.

## Discussion

The aim here was to demonstrate a high-throughput method that could be used for studying both physical interactions as well as the genetic heterogeneity in complex microbial communities by permitting high-throughput sequencing of amplicons, the origin of which can be traced back to their cellular origin by cell-specific barcodes. The method combines polyacrylamide bead encapsulation of cells with bead-localized PCR amplification of specific genomic targets and their split-pool barcoding. The method was designed to require no specialized laboratory infrastructure, such as microfluidics, to permit an easy adoption in any laboratory setting.

The lysis of the bacterial cells that took place during the polymerisation of the polyacrylamide beads was efficient enough to release sufficient amounts of DNA for the split-pool barcoding protocol. Furthermore, the polyacrylamide bead matrix was durable enough to withstand various conditions throughout the protocol, including NaOH and ether treatments, high temperatures during the PCRs and split-pool incubations as well as multiple centrifuging rounds.

In the experiment, one set of beads contained only *A. tumefaciens* cells and the other set *C. testosteroni* and *B. bullata* cells. Although some barcodes were shared between *A. tumefaciens* and *B. bullata* or *C. testosteroni*, the level of crosstalk between the beads was minor and clearly the highest proportion of barcodes were shared between *B. bullata* and *C. testosteroni*. Alongside the 16S rRNA gene, *A. tumefaciens*-specific target was included as another target for barcoding, with the aim of *A. tumefaciens*-specific target only sharing barcodes with the *A. tumefaciens* 16S rRNA gene target. Again, although some level of crosstalk was visible between the beads, the highest number of barcodes were shared between *A. tumefaciens*-specific target and the *A. tumefaciens* 16S rRNA gene target. There are multiple potential sources of crosstalk between the beads. One of them is instability of the emulsion during the emulsion PCR step, which potentially leads to the transfer of some amplicons between the beads. Another potential source of crosstalk is the template switching taking place during the PCRs performed after split-pooling.

The integrity of the barcode region varied between the 16S rRNA gene reads and the *A. tumefaciens*-specific target reads: In 72 % of the 16S rRNA gene reads the barcode region was correctly formed, whereas in the *A. tumefaciens*-specific target reads only 16 % contained all the expected barcode parts. This was also reflected in the gel electrophoresis results, where the 16S rRNA gene main product was of the correct size, but the *A. tumefaciens*-specific target amplicons were not. The low amount of correctly formed product in the *A. tumefaciens*-specific target reads was mostly explained by the lack of overhang region between the target amplicon and the first barcode oligo. Further analysis revealed that in most of the reads with no overhang, majority of the 3’ end of the *A. tumefaciens*-specific target was also missing, and the barcode sequence was directly attached to the 22-nt-long 5’ end of the target, equivalent to the forward primer sequence. Potential sources for this include secondary binding sites of the primers or the first linker, insufficient multiplex emulsion PCR or secondary structures during the PCRs ^30^. The sequence of the *A. tumefaciens*-specific target is somewhat GC-rich (approx. 62 %), which could hinder the PCR ^31^. Regardless, in majority of the reads the barcode region outside the overhang region was intact, meaning the split-pooling itself was successful.

The varying amounts of different bacterial cells used when making the beads was reflected in the results, as the strain *C. testosteroni* was the most abundant of the three strains. As the polyacrylamide beads were overloaded with bacterial cells, i.e. one bead on average contained multiple cells, the method is not yet optimised for single-cell resolution. Furthermore, all the bacterial strains used were gram-negative and belonged to the phylum Pseudomonadota. Therefore, the performance of the method with gram-positive bacteria or bacteria from other phyla remains to be tested. Additional enzymatic lysis, such as the one described by Spencer et al. (2015)^19^, might be required for diverse environmental samples.

Since the split-pool barcoding of eukaryotic cells and RNA was first described in 2017 ^21,22^, it has been utilised for microbial single-cell RNA sequencing in novel methods such as microSPLiT ^32^. Here we describe a method that combines split-pool barcoding with polyacrylamide beads for the analysis of any bacterial genomic targets that are amplifiable by PCR. The method can be utilised for the spatial analysis of microbial communities as well as linking genetic traits to single cells and has high relevance for understanding microbial community ecology and phenomena such as and the spread of antimicrobial resistance ^14,33^.

## Acknowledgements

We thank the Research Council of Finland (grants 323663, 336475 and 358202) for supporting the study.

## Supplementary Information

The supplementary information is available at Zenodo (https://doi.org/10.5281/zenodo.15737316).

## Notes

https://doi.org/10.5281/zenodo.15737316

### Competing Interest Statement

The authors have declared no competing interest.

### Summary of Updates

The bioinformatics and data analysis section of the methods has been updated to clarify the read processing steps.

